# Autonomous Wireless System for Robust and Efficient Inductive Power Transmission to Multi-Node Implants

**DOI:** 10.1101/2021.02.01.429239

**Authors:** Peilong Feng, Timothy G. Constandinou

**Author notes:** Email: {, }.

## Abstract

A number of recent and current efforts in brain machine interfaces are developing millimetre-sized wireless implants that achieve scalability in the number of recording channels by deploying a distributed ‘swarm’ of devices. This trend poses two key challenges for the wireless power transfer: (1) the system as a whole needs to provide sufficient power to all devices regardless of their position and orientation; (2) each device needs to maintain a stable supply voltage autonomously. This work proposes two novel strategies towards addressing these challenges: a scalable resonator array to enhance inductive networks; and a self-regulated power management circuit for use in each independent mm-scale wireless device. The proposed passive 2-tier resonant array is shown to achieve an 11.9% average power transfer efficiency, with ultra-low variability of 1.77% across the network.

The self-regulated power management unit then monitors and autonomously adjusts the supply voltage of each device to lie in the range between 1.7 V-1.9 V, providing both low-voltage and over-voltage protection.

## I. Introduction

A number of different wireless interface techniques have been applied to implantable medical devices for transmitting power and communicating data, such as ultrasonic [1]–[4], electromagnetic [5]–[14], optical [15]–[17]. A key objective of implementing any scheme for power transmission to an implant is to avoid the need for wires, particularly percutaneous thus reducing the risk of infection. Much of the research literature focuses on improving the power transfer efficiency whilst also ensuring biological safety (e.g. maintaining thermal dissipation to an acceptable level) [18]–[20]. These schemes are typically designed for a single (or limited) number of implanted devices, and rely on relatively precise positioning and thus good alignment between transmitter and receiver coils that themselves are relatively large [21]. This is therefore not directly applicable to a scenario with multiple, freely-position implants that all need to be simultaneously powered and are of millimetre-scale [22]–[24].

Wireless power transfer for invasive brain machine interface applications are particularly challenging where the distances and alignments between external transmitter (Tx) and implantable receiver (Rx) vary based on cerebral cortex. This results in different coupling coefficients to each of the Rx coils that in turn results in a widely variable, and unpredictable induced voltage on the secondary coil. Furthermore, this is not purely a static setup that can be calibrated, or configured after implantation, as the micromotion of brain itself (i.e. within the skull) can result is the received voltage fluctuating over time. Adjusting the Tx output power can compensate for secondary voltage fluctuations for a single Rx device. This however is not a practical solution in systems with multiple Rx devices.

This paper presents a wireless infrastructure that uses a resonator-array to enhance the inductive coupling to freely-floating implantable devices for achieving wireless information and power transfer (WIPT). The paper is organised as follows: the wireless concept is first described in Section II; the enhancement to coverage (and significantly reduced sensitivity to misalignment) is then demonstrated through comparing this to a number of typical schemes, described in Section III; Section IV then presents the self-regulated power management unit to tackle uneven spatial PTE by automatically tuning the self-resonant frequency at each secondary node (i.e. freely-floating implant).

## II. System Overview

The proposed WIPT system architecture is shown in Fig. 1. This fundamentally facilitates a *Power In & Data Out* function. The downlink is used to transmit power wirelessly from a single external transceiver to multiple freely-positioned implants through the relay resonator. In a BMI application, each of these implants will contain instrumentation circuits to record neural activity from electrodes. The uplink is then used to send data from each of these freely-positioned implants to the external device. The external device will thus collect data recorded across a distributed network of implants.

**Fig. 1.**
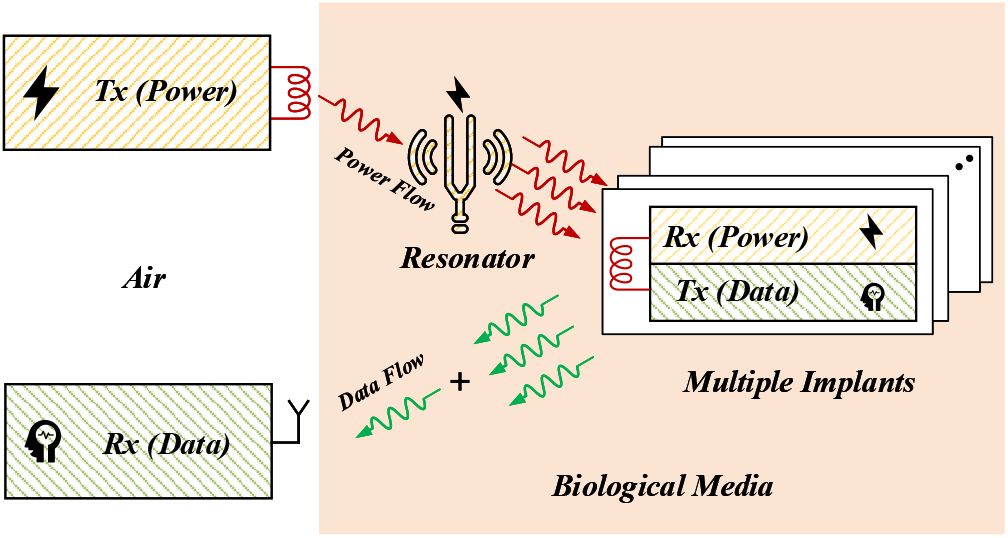
Simplified system architecture for proposed wireless information and power transmission (WIPT) scheme.

The proposed WIPT scheme provides three key features for a distributed wireless power transmission and data communication system:

1. Enhanced power transfer efficiency: The maximum inductively coupled power transmission efficiency occurs on the same resonant frequency of Tx and Rx LC tank. At the resonant condition, electromagnetic energy is coupled to the Rx LC tank through evanescent waves [25]. However, the amplitude of the evanescent waves can be enhanced by using a resonator, which results in coupling coefficient enhancement between the Tx and Rx, and eventually improves the PTE of the wireless power transfer (WPT) system based on resonant inductive coupling mechanism [6].
2. Uniform power coverage: Power coverage is determined by the magnetic flux density *B*. The resonator evenly redistributes the magnetic field density over a wider area than that of the Tx coil. Therefore, a Rx coil positioned outside the ‘coverage’ of the Tx alone can obtain sufficient power.
3. Low power consumption: The system uses back-scatter via the power link to modulate data and achieve the uplink without requiring any additional data carrier [26].

From a biological perspective, this scheme provides three key features: (1) reduced risk of infection risk due to no need for percutaneous wiring; (2) reduced tissue response and thus improved reliability due to avoiding the need for any tethers or implanted leads (e.g. [27], [28]); (3) for distributed implants, the avoidance of thermal ‘hotspots’ due to a uniformly distributed field.

## III. Wireless Power Transfer Schemes

The first step in designing a WPT system for an distributed implant network, is to select the which coil configuration is most appropriate. In this section, we compare the following 4 schemes in terms of PTE and power coverage: 2-coil, 3-coil, 3-coil with a 1-tier resonator array, and 4-coil with a 2-tier resonator array.

### A. Power Transfer Efficiency of four WPT Schemes

#### 1) 2-coil

Wireless power transmission conventionally uses a 2-coil inductive link. However, for coil pairs with a relatively large separation relative to the coil diameters, this suffers from a low power transfer efficiency due to the weak coupling coefficient between Tx and Rx coils. Furthermore, a 2-coil configuration requires near perfect alignment between Tx and Rx coil. Its power transfer efficiency is thus highly sensitive to positional and angular misalignment [29]. This is because it utilises a single Tx (primary) coil that generates a centrally focused electromagnetic field that is undesirable if requiring uniform coverage.

Fig. 2(a.1) shows a simplified equivalent schematic of 2-coil inductive link. This contains an external voltage source *V*_*s*_, source impedance *R*_*s*_ at the primary side, and load impedance *R*_*L*_ at the secondary side. According to reflected load theory [30], the PTE of a 2-coil inductive link can be expressed as

**Fig. 2.**
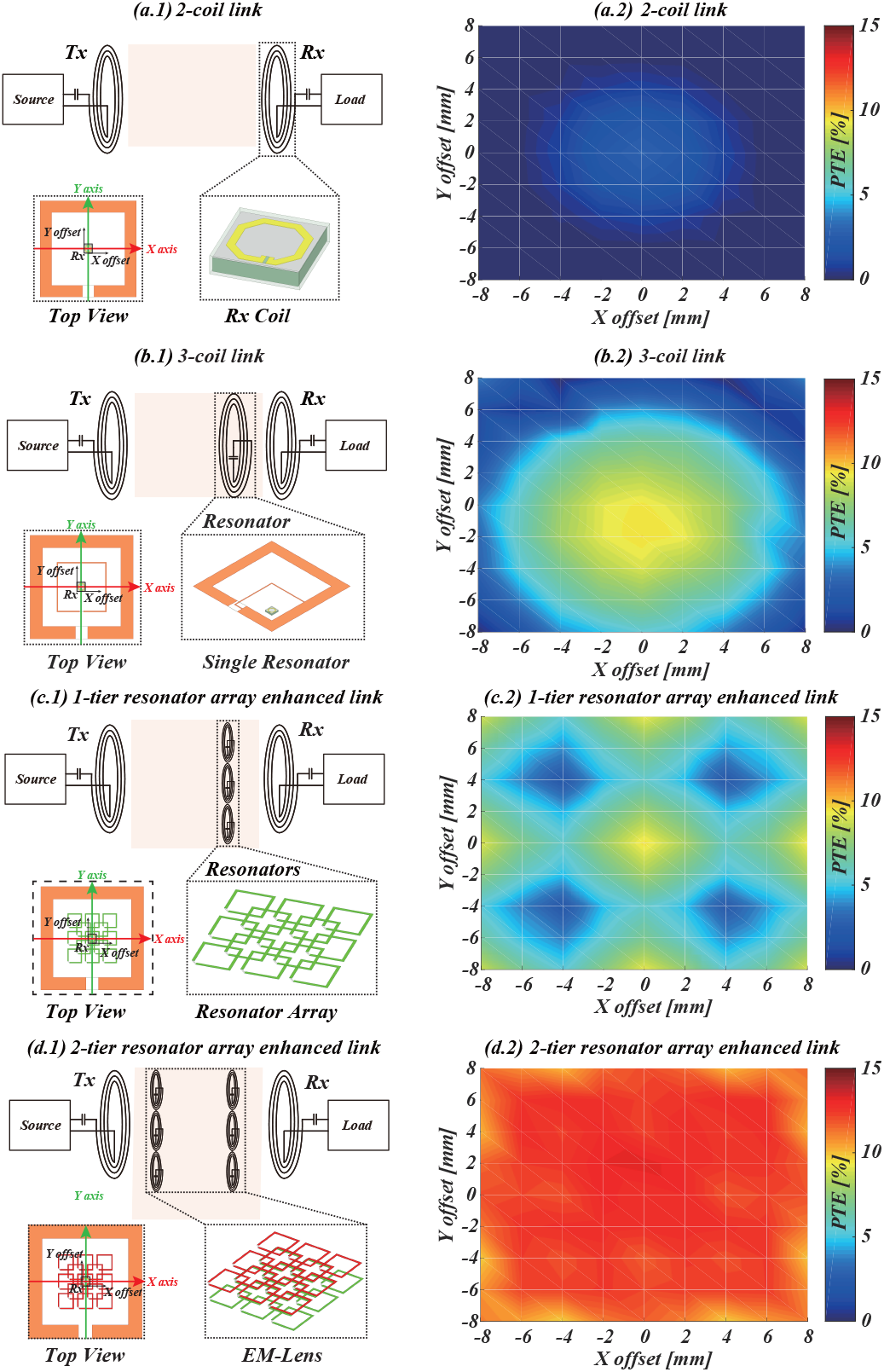
Simulation models and corresponding simulated results of power transfer efficiency across 16 mm *×* 16 mm area: (a) 2-coil; (b) 3-coil; (c) 3-coil with 1-tier resonator array; and (d) 4-coil with 2-tier resonator array.

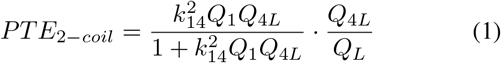

where the quality factor *Q*_1_ of the primary L_1_C_1_ tank and loaded quality factor *Q*_4*L*_ of secondary L_2_C_2_ tank are written as: *Q*_1_ = *ωL*_1_*/R*_1_, *Q*_4*L*_ = *Q*_4_*Q*_*L*_*/Q*_4_ + *Q*_*L*_, where *Q*_4_ =*ωL*_4_*/R*_4_, *Q*_*L*_ = *R*_*L*_*/ωL*_4_.

#### 2) 3-coil

The 3-coil configuration introduce a resonator *L*_3_*C*_3_ (i.e. third coil) between the primary *L*_1_ and secondary *L*_2_ coils (of the 2-coil configuration). The expression for the PTE of the 3-coil configurations consists of two terms:

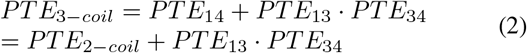

where the first term *PTE*_14_ is the same as the 2-coil link in Eq. 1, describing the PTE from the primary to secondary side. The second term *PTE*_13_ · *PTE*_34_ represents the coupling through the resonance coil. The PTE from primary coil *L*_1_ to resonator *L*_3_ is defined as [30]:

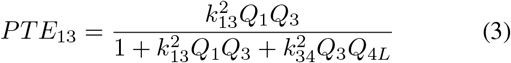

and the PTE from resonator *L*_3_ to secondary inductor *L*_4_ is defined as:

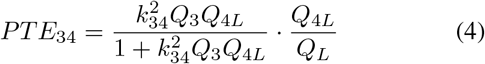

The PTE of the 3-coil link is therefore greater than that of 2-coil link (*PTE*_3*−coil*_ *> PTE*_2*−coil*_).

#### 3) 3-coil with 1-tier resonator array

Here the single resonator *L*_3_ within a simple 3-coil configuration is replaced by an array of coils *L*_3*j*_, as shown in Fig. 2(c.1). Similar to the 3-coil configuration, its PTE is also composed of two parts:

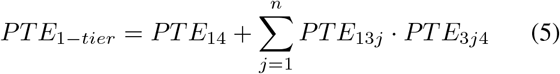

where *PTE*_13*j*_ and *PTE*_3*j*4_ represent the PTE of *L*_1_ and *L*_4_ coupled with each resonator unit *L*_3*j*_, and their equations are same as Eq. 3 and 4.

#### 4) 4-coil with 2-tier resonator array

The 2-tier resonator array implements the two intermediate coils within a 4-coil configuration using two sets of resonators *L*_2*i*_, *L*_3*j*_. The difference between this configuration and the EM-Lens previously reported in [14] is that the wired connections between each resonator unit have been removed. Also, the upper and lower resonator arrays here are independent with no connections between the two tiers. Thus, the self-resonant frequency (SRF) deviation of each resonator unit caused by internal wired connections can be eliminated, which results in PTE and coverage improvement. The PTE can here be defined as:

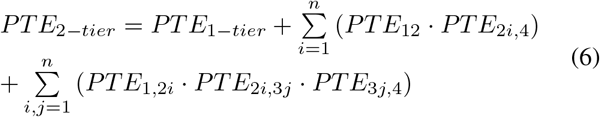

#### 5) Comparison

Comparing the expressions for PTE (1), (2), (5), (6) of the various configurations, the following order can be concluded:

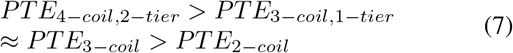

### B. Electromagnetic Simulation Verification

To verify the analytical expressions for PTEs and the comparison between the different schemes, 3D models are designed for each configuration and evaluated using a six-layer head model in HFSS (ANSYS, Canonsburg, PA). Most importantly, the spatial distribution of the PTE across the X and Y axes is an essential parameter for the design of freely positioned implants. The simulation models used for the four configurations are shown in Fig. 2, annotated with coil positions. Simulated results of spatial distribution of PTE within *±*64*mm*^2^ are included alongside each scheme.

The power coverage of the 2-coil and 3-coil configurations have a focused power profile, as shown in Fig 2(a.2,b.2). The maximum PTE is located at the central position of the Tx coil. The PTE degrades significantly as the Rx coild moves away from the centre of the Tx coil. The maximum PTE of the 3-coil link here is twice that of the 2-coil link. The average PTE of 3-coil and 2-coil link can achieve to 4.67% and 0.44%.

When the single resonant coil (i.e. in a 3-coil link) is replaced with an array of resonators, as shown in Fig. 2(c.2), the PTE can be evenly distributed over a increased range. To evaluate the PTE and uniformity of power coverage across the different schemes, the figure of merit (FoM) can be defined as: 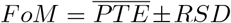 where 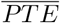 the average value of PTE is expressed as: 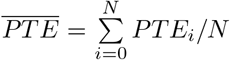. The relative standard deviation (RSD) of PTE can reveal the PTE fluctuations over a wide range. This is defined as:

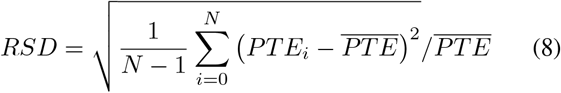

According to this expression, and the simulation results provided by HFSS, the FoM of each configuration is summarised in Table I. The essential requirements of wireless power transfer systems for multiple Rx devices are high PTE and low deviation in the spatial distribution of power. Simulation results confirm that the performance of 2-tier array significantly increases the average PTE to 11.9%, whilst reducing the power distribution deviation to 1.77%.

**TABLE I.** FoM for each inductive link.

| FoM | Average PTE | Power Distribution Deviation |
| --- | --- | --- |
| 2-Coil Link | 0.44% | 93.79% |
| 3-Coil Link | 4.67% | 13.71% |
| 1-Tier Resonator Array | 6.11% | 1.83% |
| 2-Tier Resonator Array | 11.9% | 1.77% |

## IV. Self-Regulated Wireless Power and Data

### A. System Implementation

Fig. 3 shows the system architecture of the self-regulated power management unit. This uses a Schottky diode based full-wave rectifier, supply voltage monitor integrated with a bandgap voltage reference, a digital tuning capacitor array *C*_*T*_, and finite state machine (FSM) for automatic tuning.

**Fig. 3.**
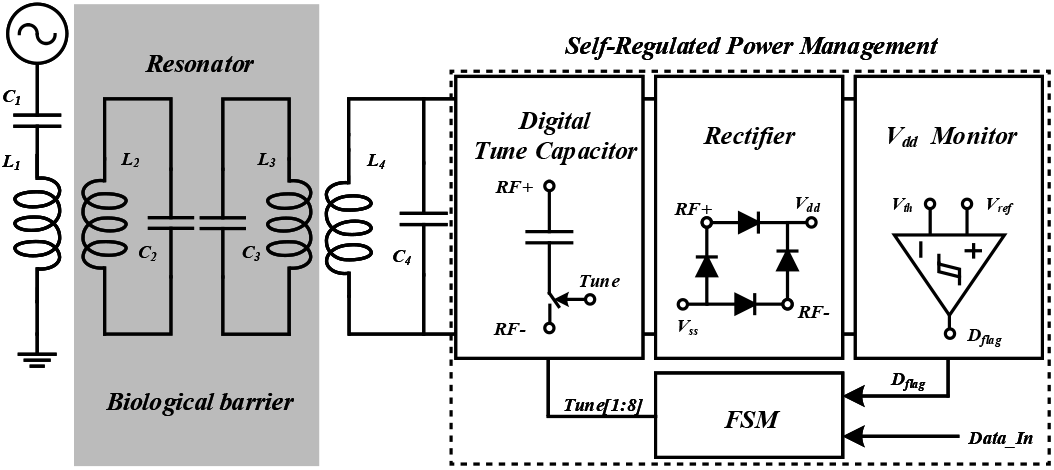
Block diagram of the self-regulated power management unit.

This system operates as follows: the AC-DC conversion is first achieved using a full-wave rectifier. The rectified DC voltage *V*_*dd*_ passes through a potential divider to define high- and low-voltage thresholds *V*_*th*_. These threshold voltages are compared with the 1.24 V bandgap reference voltage *V*_*ref*_, and generate flags *D*_*flag*_ to represent whether the supply voltage is within the specified threshold range. The finite state machine (FSM) generates 8-bit tuning data based on these flags to control corresponding bits of the capacitor array *C*_*T*_ . The tuning capacitors *C*_4_, *C*_*T*_ and the Rx coil *L*_4_ determine the self-resonant frequency (SRF) of the secondary LC tank. The voltage received on the secondary side can be effectively regulated by tuning its SRF because the entire inductive link is resonance-based. The resulting supply voltage *V*_*dd*_ is finally checked using the over- and under-voltage flags generated by the supply voltage monitor. This self-regulated power management unit therefore ensures that the DC supply voltage remains within 1.7 V - 1.9 V range.

### B. Self-Regulated Power Management

#### 1) Supply Voltage Monitor

The supply voltage monitor is designed to detect any fluctuation of the power supply voltage, and generate flags for the FSM to retune the system. This circuit is composed of a resistive divider, two strong-arm comparators [31], shown in Fig. 4. The high *V*_*dd*(*H*)_ and low supply voltage threshold *V*_*dd*(*L*)_ are defined as: *V*_*dd*(*H*)_ = *V*_*ref*_ (*R*_1_ + *R*_2_ + *R*_3_)*/R*_3_ = 1.9*V* and *V*_*dd*(*L*)_ = *V*_*ref*_ (*R*_1_ + *R*_2_ + *R*_3_)*/*(*R*_2_ + *R*_3_) = 1.7*V* . When the supply voltage *V*_*dd*_ is higher or lower than *V*_*dd*(*H*)_ or *V*_*dd*(*L*)_, the corresponding digital flags *D*_*H*_, *D*_*L*_ are set to 1. Two digital flags generated by comparators are reset to low, When supply voltage *V*_*dd*_ is adjusted within the thresholds (1.7 V - 1.9 V).

**Fig. 4.**
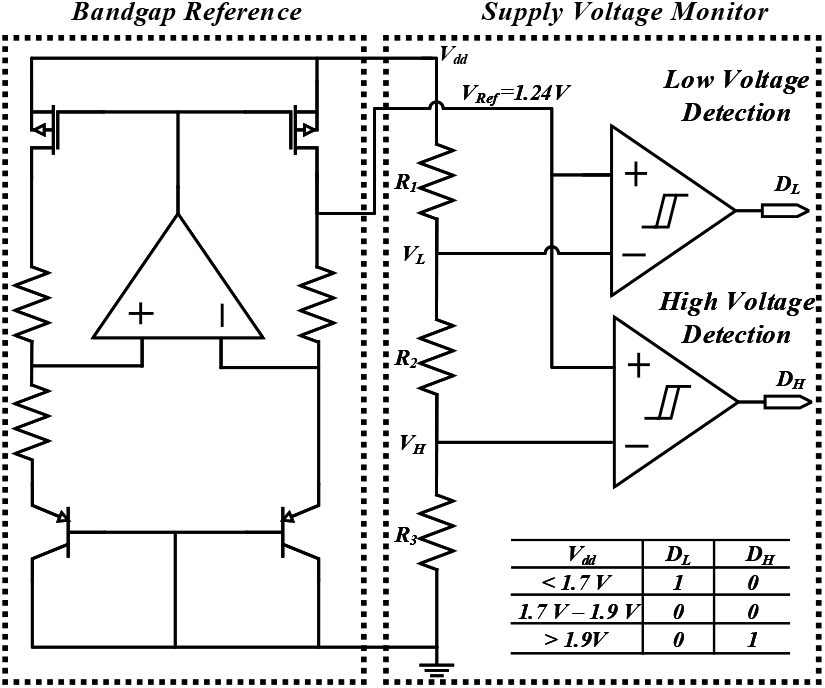
Functional diagram of proposed supply voltage monitor.

**Fig. 5.**
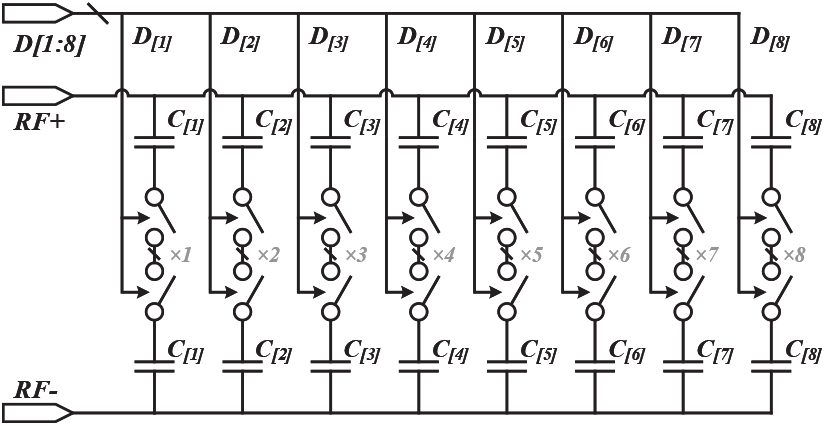
Schematic of binary weighted digital tuned capacitor array.

#### 2) Auto-Tune FSM

A simple four-state FSM is implemented to read and control the switches in the supply voltage monitor, and then to generate a 8-bit digital output for adding or removing capacitance from the secondary LC tank. The four states are idle, compare, tune and data. The FSM firstly ensure supply voltage fluctuations within the thresholds, and then the input data is transferred by modulating the power carrier through backscatter.

#### 3) Digital Tune Capacitor

The unit capacitance of the 8-bit binary-weighted capacitor array is 252.32 fF (including parasitic capacitance), with a total capacitance of 9.08 pF. The SRF is inversely proportional to the square root of capacitance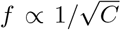. SRF variations lead to induced voltage fluctuations on the secondary side. The maximum induced voltage is achieved when the secondary SRF is the same as the relay resonator. The induced voltage curve in terms of operating frequency conforms to the standard normal distribution. When the secondary SRF is close to the primary SRF, the unit capacitance may cause significant voltage changes in the induced secondary voltage. The initial value of the binary-weighted capacitor array is set to 128 to ensure sufficient tuning range from 381.89 MHz to 509.99 MHz. According to the simulation results in Fig. 6, the rectified voltage variation is from 1.85 V to 2.31 V. The self-regulated voltage can remain within *V*_*dd*(*H*)_ and *V*_*dd*(*L*)_ thresholds.

**Fig. 6.**
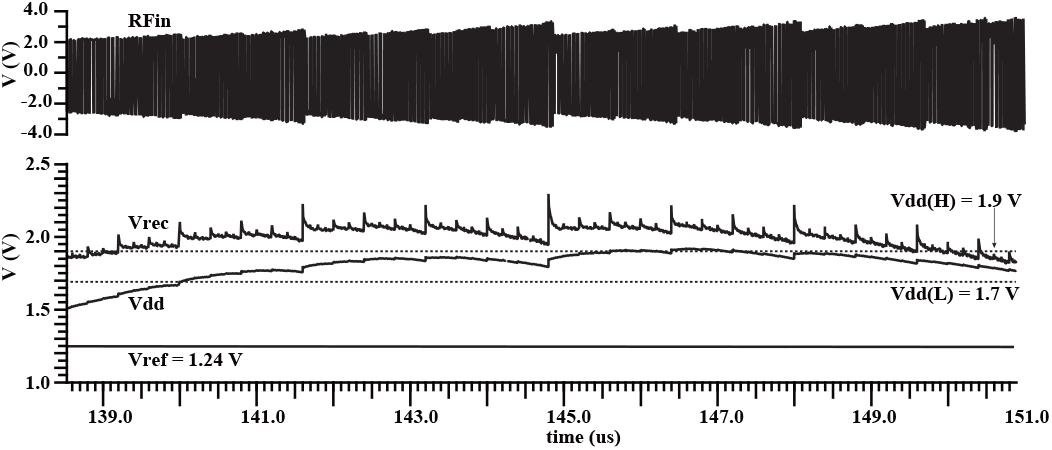
Simulated results of self-regulated power management unit.

### C. Backscatter Data Communication

Passive backscatter communication scheme consumes less power than active modulated communication schemes, such as UWB, BLE, as this passive modulation doesn’t need to generate data carrier on-site. Here we use a 500 fF capacitor to achieve 1.47% SRF offset, so that the transmitting energy can be backscattered from the secondary side to the primary side. At the same time, the voltage deviation caused on the secondary side is within 0.1 V, which will not affect the stability of the supply voltage.

## V. Conclusion and Future Work

This work has proposed 1- and 2-tier resonator coupled inductive links for simultaneously powering an arbitrary number of implant devices regardless of their specific location and/or orientation. The spatial power distribution has been analysed and compared against four different WPT schemes. This can guide the selection of a candidate WPT scheme for future implantable devices. Secondly, a circuit for self-regulating the power supply has been proposed. This can enable a distributed network of autonomous, independent implant nodes.

The proposed circuit has been designed and fabricated in a commercially-available 180 nm technology. Future work will integrate on-chip coils to form the secondary LC tank using stack die wire bonding [32]. The resonator array will be implemented using a flexible PCB substrate. The network will be validated using a software defined radio-frequency platform. This WIPT scheme will enable variety of multi-access systems to enable transmission wireless power transmission and data communication for multiple implants.

## Acknowledgement

This work was supported by the Engineering and Physical Sciences Research Council (EPSRC) grant EP/M020975/1.

